# Are alpha-band oscillations a default dynamic of the human visual brain, or a product of ongoing visual processes?

**DOI:** 10.1101/2020.02.20.958900

**Authors:** Wiremu Hohaia, Blake W. Saurels, Alan Johnston, Kielan Yarrow, Derek H. Arnold

## Abstract

One of the seminal findings of cognitive neuroscience is that the power of alpha-band (∼10 Hz) brain waves, in occipital regions, increases when people close their eyes. This has encouraged the view that alpha oscillations are a default dynamic, to which the visual brain returns in the absence of input. Accordingly, we might be unable to *increase* the power of alpha oscillations when the eyes are closed, above the level that would usually ensue. Here we report counter evidence. We used electroencephalography (EEG) to record brain activity when people had their eyes open and closed, before and after they had adapted to radial motion. The increase in the power of alpha oscillations when people closed their eyes was *enhanced* by adaptation to a broad range of radial motion speeds. This effect was greatest for 10Hz motion, but robust for other frequencies, and specifically for 7.5Hz. This last observation is important, as it rules against an ongoing entrainment of activity, at the adaptation frequency, as an explanation for our results. Instead, our data show that visual processes remain active when people close their eyes, and these can be modulated by adaptation to increase the power of alpha oscillations in occipital brain regions.

## Introduction

One of the seminal findings of cognitive neuroscience has been that alpha-band activity in visual brain regions is *enhanced* when people close their eyes (Berger, 1929; Adrian & Mathews, 1934). Long after this seminal observation, it was noted that neural spiking in cortex is *suppressed* when the power of alpha-band oscillatory activity is increased (see Haegens et al., 2011; and Lörincz et al., 2009). This encouraged the idea that *decreases* in the power of alpha-band activity, when people open their eyes, are due to a *release from inhibition* (Pfurtscheller, Stancák, & Neuper, 1996). This, in turn, prompted the suggestion that the level of alpha-band activity when people close their eyes is a *default* dynamic of the human visual brain, to which it returns in the absence of input (Pfurtscheller, Stancák, & Neuper, 1996).

We felt that, rather than an idle-like state to which the visual system returns in the absence of input, the power of alpha-band oscillations when the eyes are closed might be a product of ongoing visual processes. This speculation was informed by the fact that human eye-lids are partially transparent. If you are in a lit environment you can demonstrate this to yourself. Close your eyes, and note the apparent brightness level. Then, additionally cover your eyes with your hand. When you do so, you should note that the apparent brightness level will *further decrease*. You can make your visual world darker still, even if you have closed your eyes, by additionally covering them with your hand. This not only demonstrates that your eyelids are transparent, it also demonstrates that visual processes are *ongoing* when your eyes are closed (otherwise you would not have noticed the further darkening of the visual world when you covered your closed eyes with your hand).

As visual processing is ongoing when you close your eyes, the power of alpha-band oscillations in occipital brain regions might be a product of active visual processes, even when your eyes are closed. So, we decided to see if these could be influenced by visual adaptation. Adaptation is a ubiquitous property of sensory coding, including visual processing (Carandini & Heeger, 2011). Visual adaptation can be induced by prolonged exposure to a specific stimulus, or set of stimuli. This can alter sensory activity for a protracted period, and change perception (Webster, 2012). Such changes are observed in motion (Glasser, Tsui, Pack, & Tadin, 2011), orientation (Blakemore & Campbell, 1969), and shape perception (Storrs & Arnold, 2017) – to name just a few visual attributes. Visual adaptation is therefore a popular investigative tool for examining computational processes underlying perception, leading to its colloquial title – the psychophysicist’s microelectrode (Frisby, 1979).

If the level of alpha power, in occipital brain regions when people close their eyes, is a default dynamic to which the visual brain returns in the absence of input (Mantini et al., 2007), we might be unable to further *exaggerate* this dynamic via visual adaptation. If, however, the power of alpha-band oscillations is a product of ongoing intrinsic visual processes, even when you close your eyes, we might be able to modulate these via motion adaptation.

To explore these possibilities, we used EEG to record brain activity from occipital sensors while people had their eyes open and closed, before and after they had adapted to radial motion. We had a small number of experienced observers adapt to a range of different speeds, across multiple testing sessions, so we could describe any changes in eyes closed alpha power as a function of adapting speed. This suggested a range of effective frequencies. From these, we selected one (7.5Hz) that did not coincide with the alpha rhythm of the visual brain (∼10Hz), and tested the robustness of our core finding with a larger group of volunteer participants. We find that the power of alpha-band oscillations, recorded by occipital sensors when the eyes are closed (though true to a lesser extent for eyes-open data), can be *increased* by pre-adapting people to radial motion.

## Methods and results

### Participants

Six lab members (5 male, M age = 27.5) volunteered to participate as *experienced* observers who could be tested repeatedly. Three of these were authors (WH, BWS, DHA). To assess the robustness of findings suggested by data from these participants, we recruited a further 25 students from a research participation pool in the School of Psychology at The University of Queensland (15 male, M age = 19). All of these student participants were naïve as to the purpose of the experiment, and received course-credit in exchange for participation. All participants reported having normal, or corrected-to-normal visual acuity (we asked them to wear glasses / contact lenses if they would normally do so while reading).

### Stimuli and apparatus

Testing took place in a darkened room. A chinrest ensured a constant viewing distance (57cm). During all eyes-open periods (including adaptation), participants fixated a central red dot, with a diameter subtending 0.4 degrees of visual angle (dva) at the retina. Stimuli were presented on an ASUS VG248QE 3D Monitor (1920 x 1080 pixels, refresh rate: 60Hz), driven by a Cambridge Research Systems ViSaGe stimulus generator and custom MATLAB R2015b (The MathWorks, Natick, MA, USA) software. A Tucker-Davis Technologies Audio Workstation was used to produce audio cues, emitted diotically at supra-threshold level by speakers on either side of the testing display. The adaptor consisted of a sine-wave luminance-modulated radial grating (8 cycles; radial frequency: 4/π c/rad; contrast: 50%), presented within an annulus with an outer diameter subtending 29 dva and an inner annulus subtending 2 dva. EEG data were recorded using the Biosemi International ActiveTwo system. Electrodes (64 AG/agCI) were placed according to the extended international 10-20 system and digitised at a 1024Hz sample rate with 24-bit analog-digital conversion.

Adaptor rotation direction was unidirectional, and counterbalanced across both experienced and student participants. For experienced participants, adapting speeds (5, 7.5, 10, 15, and 18Hz) were adjusted for different blocks of trials. For these participants, blocks of trials for the different adaptors were conducted on different days, and in a pseudo random order. Student participants completed a single experimental session, adapting to 7.5Hz rotation.

### Procedure

Each experimental session began with preliminary baseline recordings (48 trials). This prevents baseline trials from being impacted by adaptation, but confounds the adaptive state with time elapsed in the experimental session. This confound was, however, constant across all adaptation frequencies, so it cannot explain any differential impact of adaptation.

On baseline trials, participants were cued to close and open their eyes, by low- and high-pitched audio tones. Subjects were given ample time (∼2.6-3.6s) to close their eyes or remain fixated. During eyes-closed periods, a uniform dark display was presented in front of the participant. During eyes-open periods, participants fixated the red dot in the middle of an otherwise grey display. In each case, recordings of brain activity were then taken for 6 seconds, after which time the participant was cued to re-open their eyes (if they had been closed). There was then a fixed ITI of 1.3 seconds before the next procedure began.

Each of 48 experimental trials in a block of trials began with the presentation of an adapting stimulus, for 15 seconds for experienced participants, and for 20 seconds for the more numerous student participants. The adapting stimulus then disappeared, and after a variable ISI (2.6 to 3.6 seconds) participants were either cued to close their eyes, or there was no cue (so participants left their eyes open, see Figure 1 for a graphic depicting the experimental protocol).

**Figure 1.**
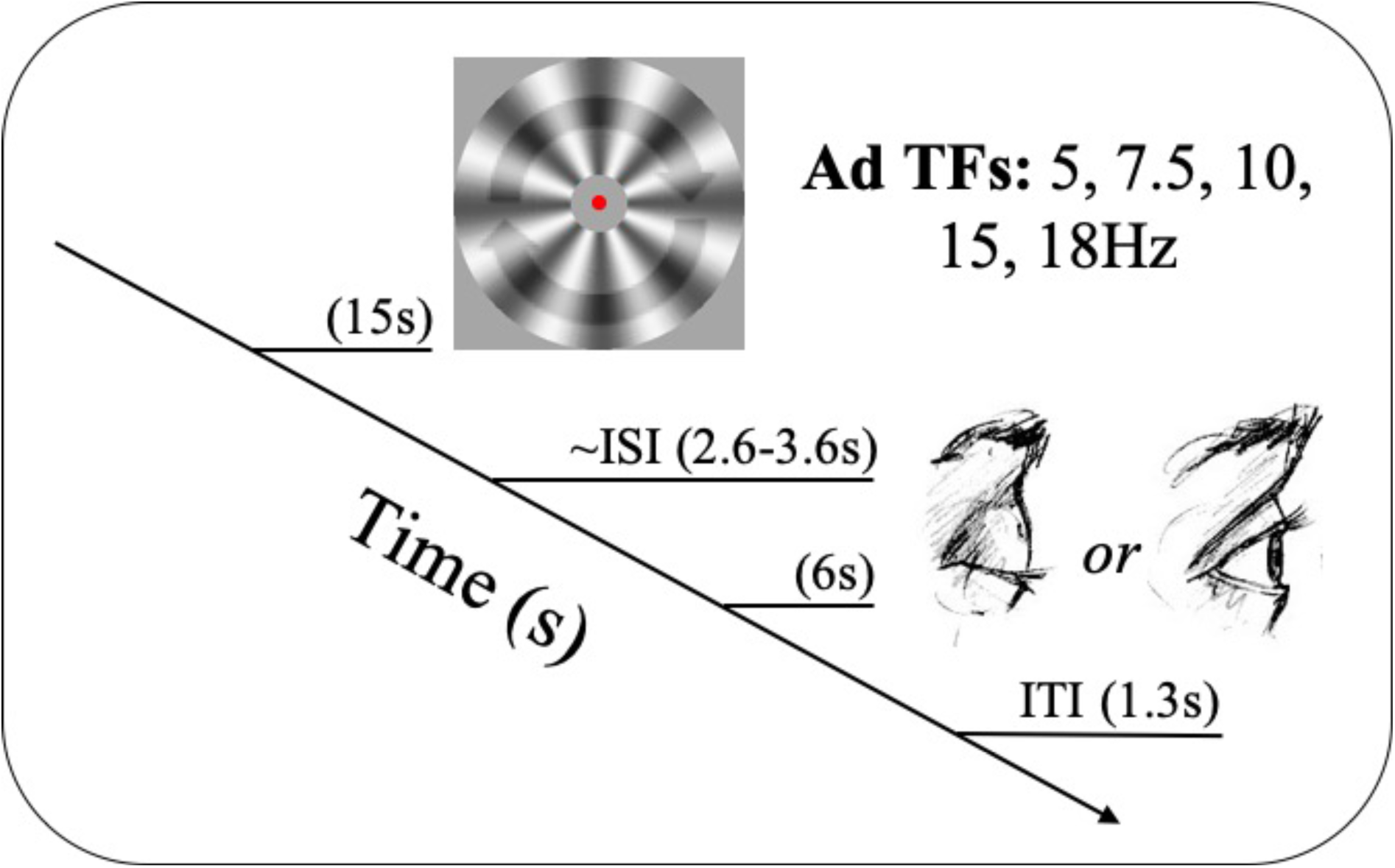
Graphic depicting the experimental protocol. Participants were adapted to radial motion for 15 seconds, animated at one of five adapting frequencies (5, 7.5, 10, 15, or 18Hz). Participants were prompted to close their eyes (by a low-pitched tone) or to keep them open while fixating centrally. A variable inter-stimulus-interval (ISI, 2.6 to 3.6 seconds) followed. Brain activity was then recorded for 6 seconds, before the participant was prompted to re-open their eyes (if they were closed) by a high-pitched tone. There was then a fixed inter-trial-interval (ITI) before the adaptor was re-presented. The procedure for baseline trials was similar, excluding adaptation periods.

### EEG data pre-processing and analyses

All analyses of data were conducted using custom MATLAB scripts involving FieldTrip toolbox (Oostenveld, Fries, Maris, & Schoffelen, 2011) and MATLAB’s in-built Fast-Fourier transform (FFT) command. All pre-processing was done offline. Data were high-pass (2Hz), low-pass (60Hz), and band-stopped filtered (49-51Hz). Blink artefacts were removed via a principal-components analysis (PCA). Data were re-referenced to volume average, and epoched into six-second segments, separately for eyes open and closed trials, for the baseline and adapted conditions.

For each participant, spectra were calculated on data from all 64 channels. Individual peak-frequency was estimated as the frequency (within 3-17Hz) at which the *largest* power difference had been measured between baseline eyes open and closed trials.

**Figure 2.**
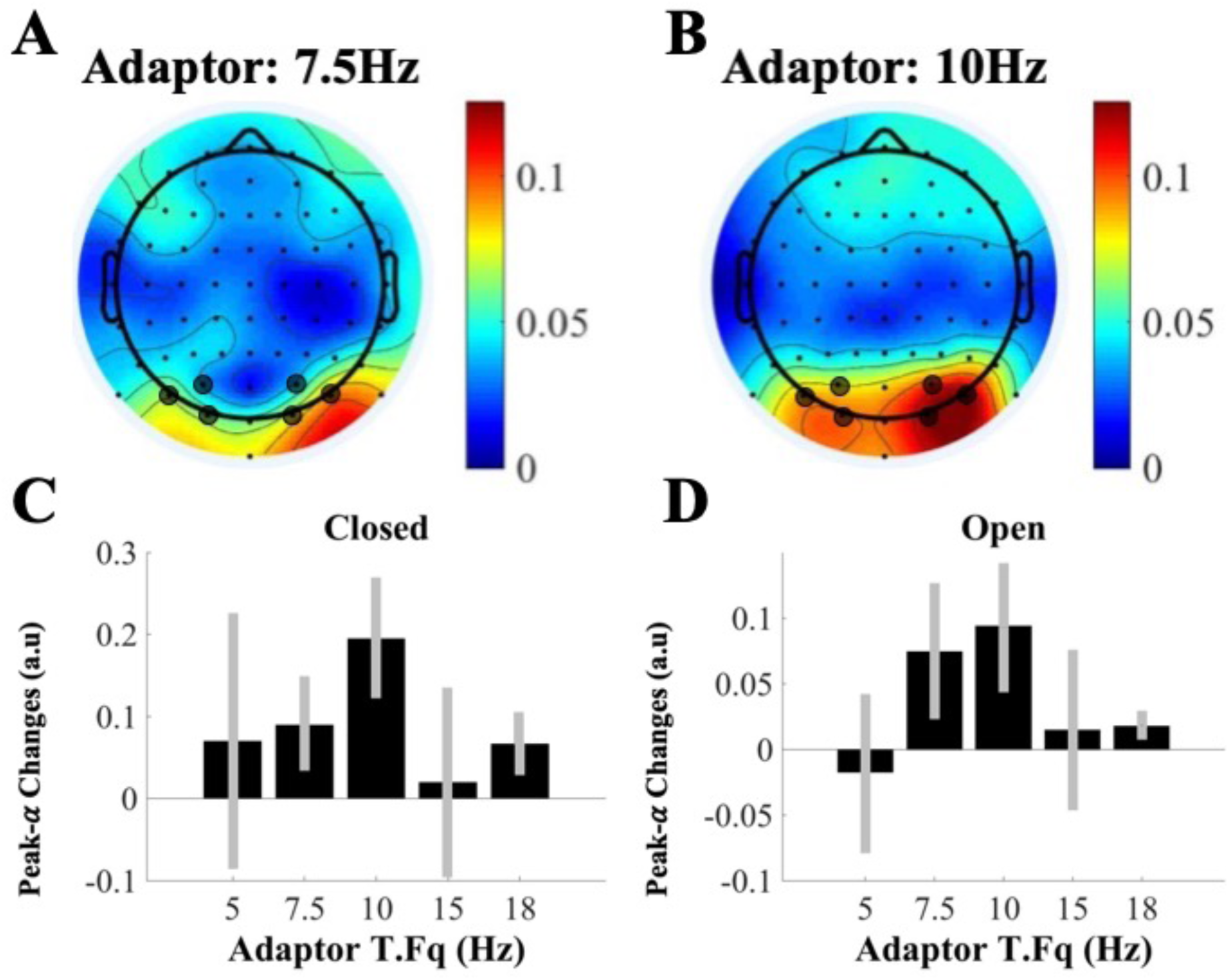
Results for experienced participants. Positive values indicate an *increase* in individual peak alpha power (∼10Hz) on adaptation trials, relative to baseline estimates. A). Topographical map of peak alpha changes when the eyes were closed, after adapting to 7.5Hz rotations. Sensors averaged to produce data depicted in Figure 2C and 2D are marked. B) As for A, but for the 10Hz adaptor. C) Peak-alpha changes (adapted – baseline) when the eyes were closed as a function of adapting temporal frequency. Error bars denote ±1SEM. D) As for C, but for eyes-open data. Note, data in C & D are plotted on dissimilar scaling.

Differences between adaptation and baseline conditions were calculated for each individual, separately for eyes open and closed trials. To illustrate the spatial distribution of adaptation-driven changes, changes in peak-alpha power at all electrodes are plotted, averaged across experienced participants. These illustrations relate to changes after adapting to 7.5Hz (Figure 2A) and to 10Hz (Figure 2B) radial motions. Note that post-adaptation, increases in alpha power seem to cluster about occipital sensors.

For experienced participants, adaptation-induced changes in alpha power recorded by occipital sensors (O1, O2, PO3, PO4, PO7, PO8) are illustrated in Figure 2C (for recordings taken when the eyes were closed) and 2D (for recordings taken while the eyes were open). These data describe a broadly tuned function, maximal for 10Hz adaptation, but also seemingly robust for 7.5Hz adaptation. Of these frequencies, we chose 7.5 Hz as the adaptation frequency for student participants, as this frequency does not coincide with average peak-alpha frequency in our sample (∼10Hz). So, using this adaptation frequency avoids a modulated frequency tag, at the adaptation frequency, from providing a credible explanation for changes in peak-alpha power.

For our student participants, both eyes-closed and eyes-open peak-alpha power was *enhanced* by 7.5Hz adaptation, relative to un-adapted baseline estimates (eyes-closed *t*_24_ = 5.62, *p* <.001, *BF*_10_ = 2442.86; eyes-open *t*_24_ = 4.12 *p* < .001, *BF*_10_ = 79.71). The change in eyes-closed alpha power was, however, *greater* than the increase in alpha power when the eyes were open (paired *t*_24_ = 4.59, *p* < .001, *BF*_10_ = 232.50; see Figure 3B).

**Figure 3.**
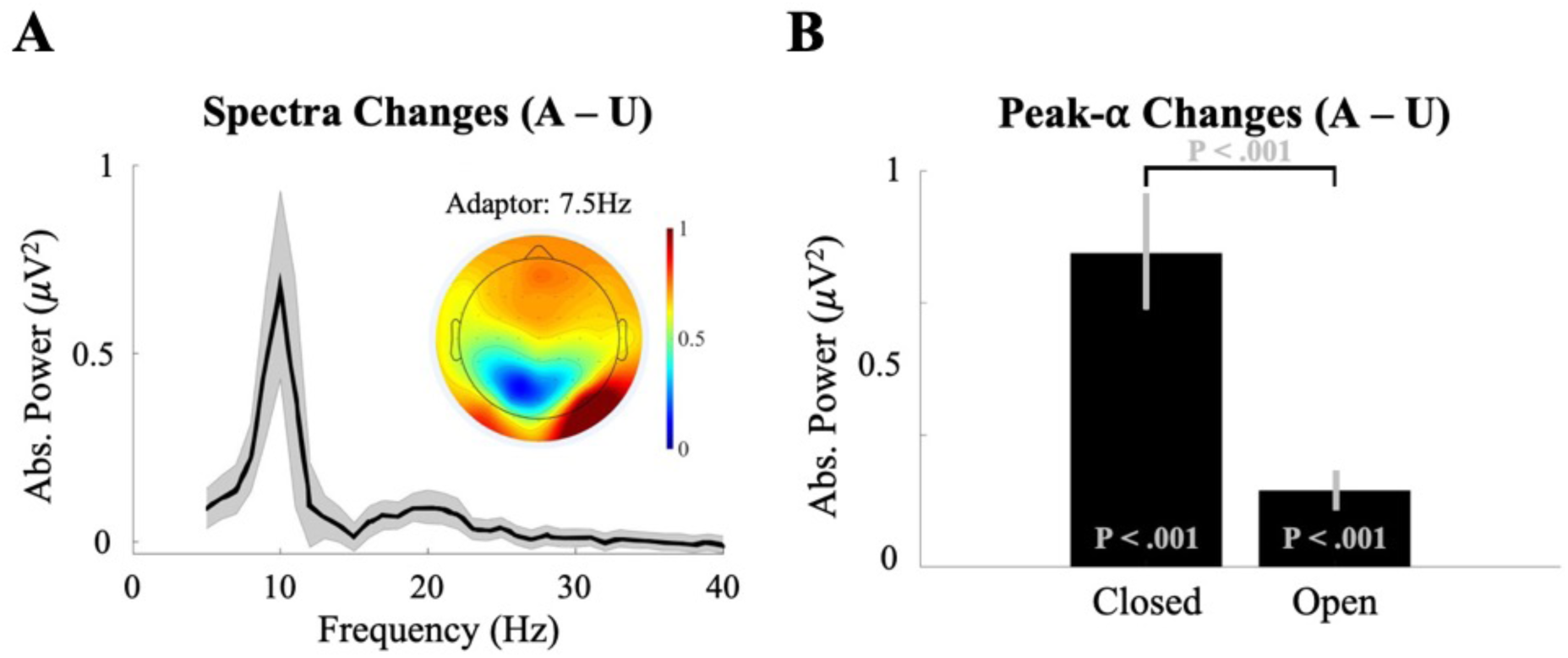
Experiment 2 results. In both plots, positive values represent an increase relative to baseline. A). Grand-averaged absolute power spectra differences, between adapted and baseline eyes-closed data as a function of frequency. Data are taken from occipital sensors: O1, O2, PO3, PO4, PO7, PO8. Shaded region depicts ±1SEM. A-inset). Group-level topographical map of adaptation driven alpha (∼10Hz) changes during eyes closed periods. B). Average individual peak-alpha power changes by condition. Data are first averaged across occipital sensors for each individual, and then across individual participants. Error bars depict ±1SEM between changes for each participant.

Due to the nature of the experimental paradigm, there is one important issue to address. To avoid any possibility of adaptation impacting baseline estimates of alpha power, adaptation always followed baseline trials. This raises the question of whether time elapsed in the experiment, or fatigue, might explain our results (see Gharagozlou et al., 2015). To address this issue, we conducted a 2 (early / late adaptation trials) × 2 (testing condition, eyes open / closed) repeated measures analysis of variance (ANOVA), sampling two subsets of adaptation data – the first and final 14 seconds of adaptation data recorded for each testing condition, comparing each of these subsets of adaptation data to the final 14 seconds of data recorded for each baseline condition. If our putative ‘adaptation’ effect had been due to fatigue, it should have been exaggerated for the final adaptation trials, relative to the initial adaptation trials. We found no evidence for this. There was a significant effect of testing condition (eyes open / closed; F_1,24_ = 28.01, *p* < .001, η^2^ = 0.54), but no effect of testing time (early / late adaptation trials, F_1,24_ = 2.17, *p* = .154, η^2^ = 0.08), or interaction between testing condition and time (F_1,24_ = 0.06, *p* = .803, η^2^ < 0.01; see Figure 4; see Discussion for further arguments against a fatigue-based explanation of our effects). There appears an upward trend toward the end of adaptation (Late) in both eyes’ conditions. However, importantly, alpha power changes for eyes-closed data early on (in adaptation) are significantly different from zero, serving as additional counter evidence for a fatigue-based explanation – adaptation-driven changes occur early on, despite the little time that has passed in the experimental session.

**Figure 4.**
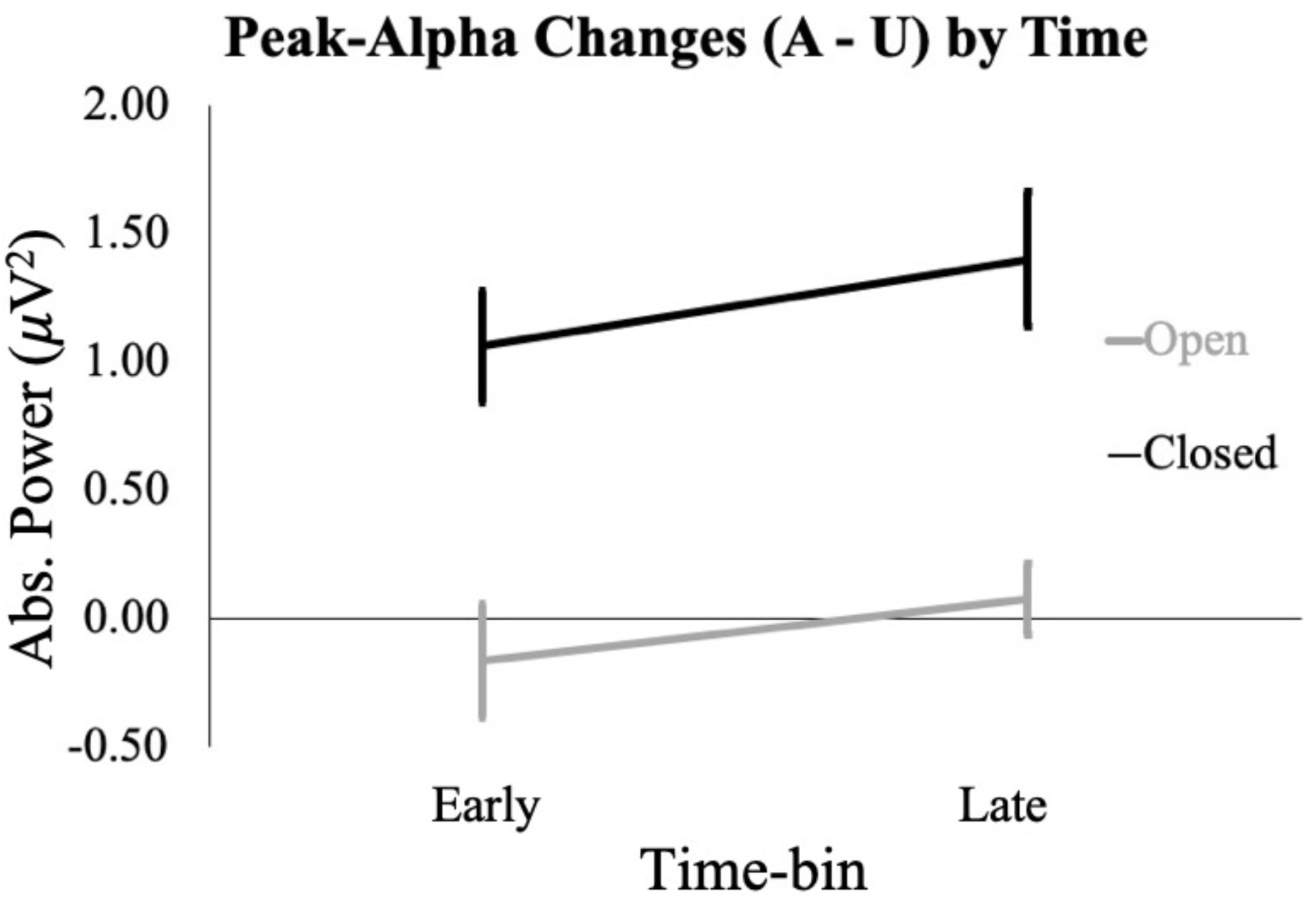
Peak-alpha changes for Early and Late adaptation periods. For these analyses, subsets of adaptation data were selected – the first (Early) and last (Late) 14 seconds of adaptation data in each experimental session, for each testing condition (Eyes Open – black, Eyes Closed – grey). Each of these are compared to the last 14 seconds of baseline data. Error bars depict ±1SEM.

## DISCUSSION

One of the seminal findings of cognitive neuroscience is that alpha-band activity in visual brain regions is *enhanced* when people close their eyes (Berger, 1929). We have shown that this effect can be *increased* by pre-adapting to radial motion. This effect appears to be broadly tuned to temporal frequencies about 10Hz, and is robust after adaptation to 7.5Hz radial motion. Peak alpha-band activity was increased by pre-adaptation to radial motion, regardless of whether the eyes were open or closed, but the adaptation-induced increase was *greater* when the eyes were closed.

To avoid contamination, adaptation always followed baseline measures. This raises the possibility that our adaptation effects might have been due to fatigue, rather than adaptation. We can discount this possibility for a number of reasons. First, our data suggest some degree of specificity, with apparent maximal adaptation effects after 10Hz adaptation. We would not expect fatigue to be selective for adaptation frequency. Second, our adaptation effects were *greater* when the eyes were closed, as opposed to open. This seems opposite to what one might expect of a fatigue-related effect (assuming closing one’s eyes is restful). Third, we found no evidence adaptation effects were exaggerated for late, as opposed to early adaptation trials. However, they were apparent immediately at the onset of the adaptation block (at least for eyes-closed conditions). Again, these findings seem counter to what one would expect of a fatigue effect. Finally, we note that the experimental task was not demanding (passive viewing of a screen), and testing sessions were brief (approximately 30 minutes), so we would not expect participants to be much fatigued by our experiment.

Long after the eyes closed effect was reported (see Berger, 1929; Adrian & Mathews, 1934), it was shown that during periods of increased alpha-band synchronisation, neural spiking in cortex is *suppressed* (see Haegens et al., 2011; and Lörincz et al., 2009). This encouraged the idea that *decreases* in alpha-band activity when people open their eyes are due to a *release from inhibition* (Pfurtscheller, Stancák, & Neuper, 1996). This, in turn, prompted the suggestion that the level of alpha-band activity when people close their eyes represents a *default* dynamic of the human visual brain, which is disrupted when people open their eyes (Pfurtscheller, Stancák, & Neuper, 1996). Our data challenge this view.

Rather than an idle-like state, to which the visual system defaults in the absence of input, our data suggest that eyes-closed alpha oscillations are, at least in part, a product of ongoing visual processes. Otherwise, they should not have been impacted by visual adaptation, which modulates visual processing (Webster, 2012). We feel the classic account of the eyes-closed alpha effect misconstrues eyes-closed conditions, as periods when all visual input is eliminated, and visual processing ceases. The eye-lids are partially transparent. Effectively, closing your eyes does not instigate a cessation of visual processing.

One interesting facet of our data is that we were able to use adaptation to 7.5Hz radial motion to *increase* peak alpha-band (∼10Hz) oscillatory power in visual brain regions. This is interesting, both because it precludes an ongoing stimulus-driven frequency tag from providing a viable interpretation of our data (Regan, 1989), and because it might provide some insight into the tuning characteristics of the mechanisms that generate the alpha rhythm. While humans can perceive a broad range of different speeds, evidence suggests this relies on mechanisms that are broadly tuned to a small number of temporal frequencies. These have been estimated as being 0Hz (static), ∼10Hz, and ∼18Hz (Hess & Snowden, 1992). It is intriguing that one of these broadly-tuned mechanisms (10Hz) would be responsive to our chosen adaptor (7.5Hz). Indeed, the broadly tuned profile suggested by our data (Experiment 1) would be consistent with this temporal-frequency tuned channel mediating the effects we have discovered.

While our data demonstrate that alpha-band oscillations in visual brain regions can be *enhanced* by pre-adapting to radial motion, they do not reveal why this particular dynamic is so marked in these brain regions, particularly when the eyes are closed. Others contend that this is because this particular frequency is a default dynamic of the human visual system, to which it returns when input ceases. Our data caution against this interpretation. They establish that the power of 10Hz oscillatory activity can be *increased* by pre-adapting to radial motion. This shows that alpha-band oscillations when the eyes are closed are, at least in part, a product of ongoing visual operations. This encourages the suggestion that the power of alpha-band oscillations might generally scale with a visual operation, which is both ongoing and can be enhanced when the eyes are closed, as opposed to open. What might this be?

One possibility is that inhibition in visual brain regions *increases* when the eyes are closed, but this effect can be further increased by pre-adapting to visual motion. Experiments involving single cell recordings and spatial attention have shown that spatially targeted inhibition of visual signalling is associated with an increase in alpha power (Luck, Girelli, McDermott, & Ford, 1997; Worden, Foxe, Wang, & Simpson, 2000; Bollimunta, Chen, Schroeder, & Ding, 2008; Bacigalupo & Luck, 2019). The power of alpha-band oscillations in visual brain regions is also enhanced when attention is directed to audio inputs (Banerjee, Molholm, Snyder, & Foxe, 2011) – consistent with a general suppression of visual processing, driven by inhibition, when visual input is less attended (for a review, see Foxe & Snyder, 2011). The increase in alpha power in visual brain regions when people close their eyes might be similar – attention might be directed to other sensory inputs, including audition, resulting in visual processing being generally suppressed via inhibition. Visual adaptation might enhance this effect, by reducing the spontaneous firing rates of a large population of visual neurons, and thereby making visual brain regions more susceptible to inhibition. Obviously, these proposals are highly speculative, and serve merely to signpost our current thinking and planning for future experiments.

The firm conclusion that can be drawn from our data is that the power of alpha-band oscillations, when the eyes are closed, is at least in part a product of visual states/processes that can be modified via adaptation. This encourages a new interpretation of one of the seminal findings of visual neuroscience (Berger, 1929; Adrian & Mathews, 1934). We suggest alpha-band oscillations in visual brain regions scale with a process that is ongoing when the eyes are closed, and we speculatively suggest inhibition of visual input as a plausible candidate.

## Acknowledgements

This research was supported by a Discovery Project Grant, funded by the Australian Research Council, awarded to D.H.A. K.Y. & A.J.

